# Demonstration of different entity of appendicitis and related causes of disease through study of cluster/outbreak: Systematic Review and Meta Analysis

**DOI:** 10.1101/628586

**Authors:** Yi-Tian Guo, Guo-Zhen Liu, Shi-Yun Tan, Yi Guo

## Abstract

**Objective:** To demonstrate different entities of appendicitis and causal association between microbiota and different types of appendicitis through studying cluster/outbreak, and providing guidance to find new cluster/outbreak of appendicitis and the epidemiological evidences of infectious etiology of appendicitis.

**Data Sources:** PubMed, Embase, CNKI, WanFang, VIP, CBM from their establishment to Jan, 2019, and the references lists from retrieved reports.

**Study Eligibility:** Reports on cluster/outbreak of appendicitis and reports of case series occurring in cluster/outbreak worldwide according to CDC’s definition of cluster/outbreak.

**Data Extraction and Synthesis:** Two researchers independently assessed report quality and extracted data according to Moose. We used random effect model for meta-analysis by Meta-Analyst ß3.13 software. Study-level assessment was conducted according to investigation methods introduced by Reingold and outcome-level assessment by GRADE system. We selected outcome measures before data collection began.

**Results:** We included 10 clusters/outbreaks of appendicitis from China and USA with total 626 patients. We demonstrated two entities, type 1 appendicitis (455 patients) and type 2 appendicitis (151patients). 20 patients left were unclassified type. For type 1 appendicitis, Natural history showed progression from a non-perforated appendicitis to perforated appendicitis as described traditionally. More than 88% of patients had elevated body temperature, WBC and neutrophil percentage. For type 2 appendicitis, natural history showed that only a few patients developed into phlegmonous appendicitis (6.9%,) or acute gangrenous appendicitis (1.4%) and no perforation or periappendicular abscess. More than 78% of patients had normal body temperature, WBC and NP. The patients’ time of type 1 appendicitis is shorter than that of type 2 appendicitis. Type 2 appendicitis had different histological features from type 1 appendicitis and was associated with fusobacteria. 9 of 10 cluster/outbreak occurred in group living unity such as school and camps, and many of them showed features of infectious diseases. The bodies of evidence were high quality in Meta analysis.

**Conclusion:** Cluster/outbreak of appendicitis is more often than expected worldwide and occurred in group living unity. Sporadic perforated appendicitis and non-perforated appendicitis may be not two different entities, but different stages of a same entity, which is inconsistent with modern classification of appendicitis. Type 2 appendicitis is a new entities. Studying cluster/outbreak is a new method in finding of new entity and causal association between microbiota and different types of appendicitis. Epidemiological evidence supported infectious etiology of appendicitis.

## Introduction

Acute appendicitis has been considered as a non-communicable disease whose public health impact is underestimated. In USA, mortality rate of appendicitis (0⋅08/10^6^) is higher than that of acute respiratory disease (0⋅04/10^6^) and influenza(0⋅03/10^6^).^1^ Natural history of acute appendicitis has traditionally been believed to often progress from an non-perforated appendicitis to perforated appendicitis,^2,3^ while a new hypothesis has been proposed that perforated appendicitis and non-perforated appendicitis may be different entities with different natural history from analysis of secular trend and clinical data^4–7^, which has become modern classification of appendicitis^8^. Differential diagnosis and management for perforated appendicitis and non-perforated appendicitis are current hot topic.^9–34^ However, all these understandings of appendicitis comes from study of sporadic patients, which may results in bias of misclassification, namely can not confirm whether or not perforated appendicitis and non-perforated appendicitis are different entities or different stage of same entity. In addition, analysis of secular trend is difficult to obtain reliable conclusion because of confounding bias. Therefore study of cluster/outbreak is helpful in these regards.

Cluster/outbreak is often feature of infectious diseases. Regarding clustering of appendicitis in USA, 1984, The Centers for Disease Control (CDC) stated that the cluster offered a unique opportunity to identify possible risk factors and to search for precipitating infectious agents, and encouraged reporting such cluster/outbreak to CDC.^35–36^ Since then, no typical cluster of appendicitis has occurred until 1997. In 1997, we found a cluster of appendicitis among students at a high school in China.^37^ In 2012, Fusobacteria were also found in these clustering patients.^38^ Since beginning of 2005, we have looked for new cluster/outbreak of appendicitis. We found that clusters/outbreaks occurred in many provinces of China and were reported in English and Chinese medical journals.^37, 39–47^ However, Nobody summarized features of distribution of cluster/outbreak of appendicitis and tried to demonstrate existence of perforated appendicitis and non-perforated appendicitis, and epidemiological evidence of infectious etiology through outbreak/cluster.

The aim of this study was to provide a new method to demonstrate different entities of appendicitis and causal association between between microbiota and different types of appendicitis and to improve modern understanding from sporadic patients. A second aim was to confirm common settings of outbreak/cluster of appendicitis and to provide guidance to find new clusters/outbreaks of appendicitis worldwide. A third aim is to provide the epidemiological evidences of infectious etiology of appendicitis.

## Methods

### Data sources and search strategy

We searched PubMed, Embase and Chinese databases: the Chinese Database of National Knowledge Infrastructure (CNKI), WanFang Data, VIP Chinese Periodical Database and Chinese Biomedical Database (CBM) including academic degree thesis and dissertation, conference proceedings for studies on cluster/outbreaks of acute appendicitis. We also searched the references lists from retrieved reports to identify additional reports by hand searching. Our search included all reports of cluster/outbreak from their establishment to Jan, 2019 with no language restriction. Except English papers, no real cluster/outbreak of appendicitis was published in non-English medical journal in Pubmed and Embase. We used the following keywords treated as title/abstract to identify relevant articles in English electronic databases: appendicitis (ti, ab) AND ((cluster (ti, ab) OR outbreak (ti, ab)); In Chinese electronic databases: appendicitis (ti, ab) AND ((cluster (ti, ab) OR outbreak (ti, ab) OR school (ti, ab) OR student (ti, ab) OR troops (ti, ab) OR training (ti, ab)), supplement 1 (search strategy for Pubmed). Because most Chinese surgeons do not have awareness of cluster/outbreak of appendicitis and there were no “cluster” or “outbreak” in their reports. We add “school”, “student”, “troops” and “training” as key words to extend the scope of literature search.

### Study eligibility

We included reports on cluster/outbreak of acute appendicitis and reports of case series occurring in cluster/outbreak according to CDC’s definition of cluster/outbreak,^48^ see supplement 2. These reports of cluster/outbreak must present histological diagnosis.

When several reports were available for the same study team, we retained the latest one for analysis. If single report did not provide enough necessary information, we combined several reports from the same study team.

### Study exclusion criteria

1. Reports with no data of body temperature, WBC, NP and no results of histological examination.
2. Reports of patients’ number less than 10 during period of cluster/outbreak.
3. Reports of cluster/outbreak which were defined through increase of incidence rate of appendicitis using statistic analysis.^49–51^

### Data extraction

Two researchers (Guo YT and Guo Y) independently retrieved all eligible reports. Any disagreements were solved by discussion with the third authors(Tang SY). Data extraction table included: author, year, settings and the outcome measure introduced as follows and as presented in table 1.

**Table 1.** Epidemiologic, clinical and histological data from cluster/outbreak of appendicitis in China

| Studies | Settings | Epidemiological data | Clinical data | Histological data |
| --- | --- | --- | --- | --- |
| Martin<sup>35</sup><br>1985<br>outbreak 1 | A small Texas town of 8000 residents, USA | During the 3 month period between February and April 1984, 13 patients with appendicitis occurred. Gender:10 male and 3 female patients. The median age:13years. 10 in school age boys.<br>Transmission route: food-borne transmission. | Patients' time (Onset to operation interval) :14hours to 4 days.<br>Average patients' time: not presented. | 31% (4 of 13) patients had perforated appendicitis. The other types of appendicitis were not presented. |
| Xiao<sup>39</sup><br>1991<br>outbreak 2-3 | A middle school for outbreak 2 and a college for outbreak 3, Jilin city, Jilin Province | Total 51 patients occurred in outbreak 2 and outbreak 3.<br>Outbreak 2: 36 students with acute appendicitis, 19 students with gastroenteritis after eating overnight food in breakfast in dining hall.<br>Outbreak 3: 15 students with acute appendicitis and 24 students with gastroenteritis after eating food left for 24 hours in dining hall.<br>Total patients including acute appendicitis and gastroenteritis' ages: 16~20 years. The authors did not provide date of onset.<br>Transmission route: food-borne transmission. | 39 of 51 patients with acute appendicitis' body temperature: 37.3°C~38.5°C; No bloody purulent stool and tenesmus.<br>Patients' time : 3 h~12 h after meals.<br>Average patients' time: not presented. | 46 patients had phlegmonous appendicitis; 5 patients catarrhal appendicitis; Gross examination: 26 had stink pus among these patients. |
| Yang<sup>40</sup><br>1991<br>outbreak 4 | University, Lanzhou City, Gansu Province | Since 1950, the incidence rate has been increased for appendicitis at Northwest University for Nationalities. 59 had acute appendicitis, of 1730 Ethnic students who were enrolled from 1981 to 1985. Xinjiang Uyghur students: 4.6% (13/282) ; Tibetan students: 4.6% (12/263); Kazak students: 4.2% (3/72); Mongolian students: 3.8% (10/263); Hui students: 2.5% (13/527); the other ethnic students: 2.5% (8/323) . However, native students' incidence rate for acute appendicitis is 0.2% (17/7100) at another university simultaneously in the same city (p<0.05).<br>Transmission route: not presented. | Proportion of patients in each grade:<br>Grade 1: 50.8% (30 patients);<br>Grade 2: 30.5% (18 patients);<br>Grade 3: 15.3% ( 9 patients);<br>Grade 4: 3.4% (2 patients).<br>It indicated that new students were more susceptible.<br>Patients' time : not presented. | 17 ethnic students had acute simple appendicitis. 23 phlegmonous appendicitis and 19 gangrenous appendicitis. Among these 19 patients, 11 had peritonitis caused by perforation.*<br>*When calculating numbers of gangrenous appendicitis and peritonitis in results, we separated "19 gangrenous appendicitis" into 8 gangrenous appendicitis and 11 peritonitis, see results. |
| Guo<sup>37</sup><br>2004<br>outbreak 5 | Middle school,<br>Wuhan City,<br>Hubei Province | Cluster phase: From April 10, 1997 to June 11, 1997, 11 patients occurred at a high school, with 3.8% (10 /290 ) students .<br>Post cluster: From the end of the initial cluster until June, 2000, 20 additional outbreaks were encountered. Female patients (6.5%) are more frequent than male patients (1.9%) . All students were from remote area and isolated areas. Cluster has not occurred since 2000 because only native students were enrolled.<br>New students were more susceptible.<br>We analyzed 29 patients operated on in collaborating hospital outcome measures.<br>Transmission route: fomite transmission.<br>Pathogen: Fusobacterium. | Among total 29 patients operated on in collaborating hospital, 4 patients' body temperatures: more than 37°C (37.1°C, 37.3°C, 37.8°C, and 37.8°C). 6 patients' WBC: $11.3 \times 10^9/L \sim 12.5 \times 10^9/L$. 12 patients' NP: 72% ∼85%.<br>Patients' time : about 50 hours. | Of 29 patients examined pathologically, 2 were diagnosed as phlegmonous appendicitis and 27 as acute simple appendicitis. Histological examination exhibited diffuse or focal hemorrhages and infiltration by eosinophils and by lymphocytes in the lamina propria or within hyperplastic lymphoid follicles,. |
| Lu<sup>41</sup><br>2005<br>outbreak 6 | Camp,<br>Lanzhou City,<br>Gansu Province | During 8 week period of military drill from 1997 to 2001(Authors did not introduce month and date), 108 new soldiers (103 male and 5 female) with acute appendicitis.<br>Transmission route: not presented. | Patients had elevated WBC and elevated NP in differential WBC.<sup>#</sup><br>Patients' time: 3h∼72h.<br>Average patients' time: not presented.<br><sup>#</sup>We imputed average WBC and average NP,and average patients' time based on reference,see supplement 2. | acute simple appendicitis and acute phlegmonous appendicitis. The authors did not report proportion of the two types of appendicitis, but patients' elevated WBC and elevated NP in differential WBC suggested that new soldiers suffer from type 1 appendicitis. |
| Lin<sup>42</sup><br>2005<br>outbreak 7 | Camp,<br>Dalian City,<br>Liaoning Province | In 2004, at a camp, 7 of 52 (13.5%) Tibetan soldiers suffered from acute appendicitis and non-Tibetan soldiers 13 of 512 (2.5%), which was significantly different ($\chi^2=13.4$, $p<0.01$).<br>Transmission route: not presented. | Clinical data: not presented.<br>Patients' time: not presented. | The authors only indicated that diagnosis for appendicitis was proven by surgery and pathology, but not indicated types of appendicitis. |
| Yan<sup>43,44</sup><br>2008<br>outbreak 8 | Camp,<br>Tibet | During period of training new recruit soldiers (from Dec. each year to March, the next year) From 2004~2007, 189(175 male and 14 female) with appendicitis; Ages: 16 years~20 years, mean age: 18.8 years.<br>Transmission route: not presented. | 35 patients' body temperature: 36.5℃~37.2℃; 116 patients' temperature: 37.3℃~38.0℃; 38 patients' temperature>38℃. All patients' WBC :12×10<sup>9</sup>/L~20×10<sup>9</sup>/L and 24 patients of them >20×10<sup>9</sup>/L; All patients' NP: 72%~93%.<br>Patients' time: For most patients 2h~72 h and for 15 more patients >72 h.<br>Average patients' time: not presented. | 97 patients had acute simple appendicitis; 68 patients phlegmonous appendicitis; 24 patients perforation. Among these patients, 2 periappendicular abscess. |
| Li<sup>45,46</sup><br>2008<br>outbreak 9 | Middle school,<br>Qingdao city,<br>Shandong Province | 660 Xinjiang Ethnic students were enrolled from 2002 to 2007; 35(10 male and 25 female) students with acute appendicitis; Age: 16~18 years. The incidence rate for Ethnic students each year is much higher than that for native students and total relative risk was 89.36 for 2002-2007.<br>Transmission route: not presented. | The patients' body temperature: 37.1℃~39.3℃. All patients' NP: 71%~89%. The authors did not provide the proportion of patients with elevated WBC.<br>Patients' time: 1 h~11 h.<br>Average patients' time: 5h. | 11 patients had acute simple appendicitis; 15 patients phlegmonous appendicitis; 3 patients gangrenous appendicitis; 6 patients chronic appendicitis. |
| Guo<sup>47</sup><br>2017<br>outbreak 10 | Middle school,<br>Nanchang City,<br>Jiangxi Province | Cluster phase: From 2000 to 2010, 120 patients occurred among Tibetan students at the middle school, with female preponderance in appendicitis patients (female 20.4%, 102 of 499; male 3.8%, 18 of 474; chi-square=62.280, $P\leq 0.001$). outbreak of appendicitis never occurred among more than 7000 native students at this school.<br>We analyzed 116 patients operated on in collaborating hospital for histological examination and 117 patients for body temperature, WBC and NP.<br>Transmission route: fomite transmission. | Among total 117 patients operated on in collaborating hospital, 12 patients' body temperatures: more than 37℃. 19 patients' WBC: 10.3×10<sup>9</sup>/L~18.3×10<sup>9</sup>/L. 12 patients' NP: 71%~91%.<br>Average patients' time was about 4.7days | Of 116 patients examined pathologically, 8 were diagnosed as phlegmonous appendicitis and 2 as gangrenous, and the the rest as acute simple appendicitis. Like outbreak 5, histological examination exhibited diffuse or focal hemorrhages and infiltration by eosinophils and by lymphocytes in the lamina propria or within hyperplastic lymphoid follicles,. |

### Outcome measures

We recorded epidemiological and clinical outcomes from reports of cluster/outbreak of appendicitis and case series occurring in cluster/outbreak. The epidemiological outcome measures were data of settings and incidence percentage (or attack rate or risk), such as number of new patients with appendicitis and population among a specific group of persons during period of cluster/outbreak. The clinical outcome measures were histological examination for specimen of appendices, body temperature, white blood cell count (WBC), neutrophil percentage (NP), interval between onset of symptom to hospital (patient’s time) and test of infectious agent. Imputation of partial missing data was introduced in supplement 3. ^9,15,17,53^

We were not successful in contacting authors for the detailed information of primary study because affiliation of authors of 7 out of 9 reports except our report was military hospital, they refused to provide more data for secret reasons. The rest one did not provide exact affiliation.^40^ Data from our manuscript of primary research prepared for submission was also supplemented into this systematic review and meta-analysis.^47^

### Demonstration of natural history of appendicitis

Cluster/outbreak often results from common cause, therefore almost every patient belongs to the same entity and appeared in different stage of same entity. Connecting each stage, we can describe full natural histories of different entities and accordingly demonstrate whether or not perforated appendicitis and non-perforated appendicitis are different entities or different stage of same entity better than through study of sporadic appendicitis.^4–7^

### Report quality assessment

We used both study-level assessment and outcome-level assessment for report quality.^52–53^ Study-level assessment was conducted according to investigation methods introduced by Reingold^54^ and outcome-level assessment according to five reasons of the grading of recommendations assessment, development, and evaluation (GRADE) system,^55, 56^ namely bias of risk, inconsistency, indirectness, imprecision of results, and publication bias. GRADE specified four categories for the quality of a body of evidence for each outcome as high, moderate, low, or very low^57^, see supplement 4.

### Subgroup analysis

We classified patients from cluster/outbreak into two subgroups according their clinical features. For subgroup 1, the majority of patients have low grade fever(≈38° C), elevated WBC count, elevated NP and phlegmonous appendicitis or more severe appendicitis in accordance with description of Sabiston Texbook of Surgery.^58^ We defined these patients as type 1 appendicitis. For subgroup 2, the majority of patients had normal temperature, WBC count, NP and acute simple appendicitis, which were different from description of Sabiston Texbook of Surgery. We defined these patients as type 2 appendicitis.

### Sensitivity analysis

To judge the stability, we performed sensitivity analysis with sequential deletion of one report and compared the results before and after the deletion.^59^

### Publication bias

According to recommendations from Cochrane collaboration, We did not conduct testing for funnel plot asymmetry because less than 10 clusters/outbreaks for each outcome measure of Meta analysis.^60^

### Statistical analysis

We used equation “n=Z^2^p(1-p)/L^2”^ compute optimal information size for meta analysis in stead of the online calculator. ^61–62^ Z was 1.96. p was the percentage of the population having a particular feature. L denoted the margin of error which was set for 0.05. According to calculation, the biggest optimal information size is 378 for results of histological examination, body temperature, WBC and NP. The number of our patients met optimal information size.

We did meta-analysis using Meta-Analyst ß3.13 soft ware. I^2^ statistic was used to evaluate statistical heterogeneity of the reports included. A rough guide to interpretation from Cochrane Handbook for Systematic Reviews of Interventions Version 5.1.0 are as follows: 0% to 40%: might not be important; 30% to 60%: may represent moderate heterogeneity; 50% to 90%: may represent substantial heterogeneity, 75% to 100%: considerable heterogeneity.

Regardless of heterogeneity or not, we used to random effect model to provide a conservative estimate of the results. We conducted subgroup analysis to compare the features of subgroup 1 and subgroup 2. We compute incidence rate of appendicitis, the overall percentage of patients with phlemoneous appendicitis group, and the overall percentage of patients with elevated body temperature, WBC counts and NP, and presented the results in forest plots. In our data, Phlemoneous appendicitis group included more severe histological change, such as gangrenous appendicitis, and perforation and so forth.

## Results

### Results from analysis of reports of cluster/outbreak

Our search yielded 483 reports. After removing duplicates and read titles and abstracts, We identified 23 full-text reports of cluster/outbreak of appendicitis assessed for eligibility and finally included 9 reports and one of our manuscript of primary research prepared for submission for systematic review and meta analysis,^37, 39–47^ Fig.1. Total 626 patients’ main epidemiological data, clinical data and histological data were listed in table 1. These clusters/outbreaks occurred in 7 provinces and autonomous regions in China, and one occurring in USA.

**Fig 1.**
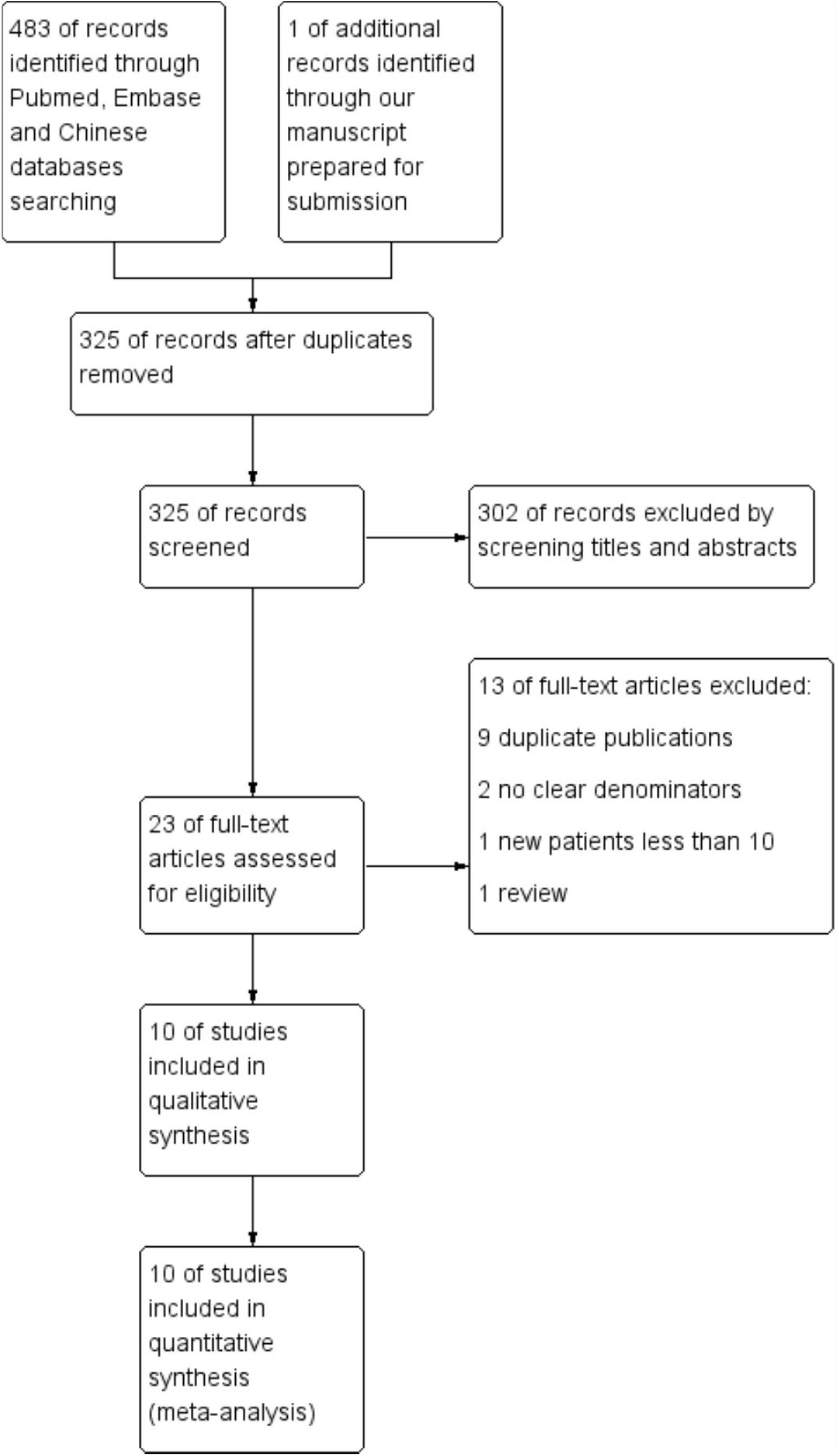
Flow diagram for selection of studies of cluster/outbreak of appendicitis in meta-analysis

Subgroup analysis demonstrated that patients in cluster/outbreak had two entity of appendicitis, namely type 1 appendicitis and type 2 appendicitis. They had different natural history and features. 20 patients left were unclassified type.

The natural history and clinical features of type 1 appendicitis (of 455 patients totally, 334 had histological examination). The patients’ natural history showed that most patients developed from classic acute simple appendicitis (37.4%, 125/334, data from outbreak 4, outbreak 8 and outbreak 9) into phlegmonous appendicitis (45.5%, 152/334, data from outbreak 2-3, outbreak 4, outbreak 8 and outbreak 9) and some eventually acute gangrenous appendicitis (3.3%, 11/334, data from outbreak 4 and outbreak 9), or perforation or periappendicular abscess or peritonitis (11.2%, 39/347, data from outbreak 1, outbreak 4 and outbreak 8). 3.3% (11/334, data from outbreak 2-3 and outbreak 9) were chronic appendicitis and catarrhal appendicitis. More than 82% of patients had elevated body temperature, elevated WBC and elevated NP in differential WBC. The above features were in accordance to description of Sabiston Textbook of Surgery. Although outbreak 1 did not present percentage of histological diagnosis, 31% (4/13) patients had perforated appendicitis at time of surgery; although outbreak 6 did not present percentage of patients with histological diagnosis, they had elevated WBC and NP in differential WBC. According to description of appendicitis of Sabiston Texbook of Surgery, the patients occurring in outbreak 1 and outbreak 6 should belong to type 1 appendicitis.

The natural history and clinical features of type 2 appendicitis (of 151 patients totally, 145 had histological examination). Appendical tissues of 145 patients were examined using immunohistochemistry. For outbreak 5 and outbreak 10, the patients’ natural history showed that most patients had nonclassic acute simple appendicitis (91.7%, 133/145; data from outbreak 5 and outbreak 10). Only a few patients developed into phlegmonous appendicitis (6.9%, 10/145) or acute gangrenous appendicitis (1.4%, 2/145) and no perforation or periappendicular abscess. More than 78% of patients had normal body temperature, WBC and NP and main histological features of appendice revealed hemorrhage and infiltration of eosinophils, which were different from type 1 appendicitis.

The forest plots of comparison for outcome measures showed obvious differences between two types of appendicitis, fig 2 (supplement 5). Among the patients with type 1 appendicitis, the percentages of elevated body temperature, WBC, NP and phlegmonous or more severe appendicitis were much higher than that among patients with type 2 appendicitis.

**Fig 2 A.**
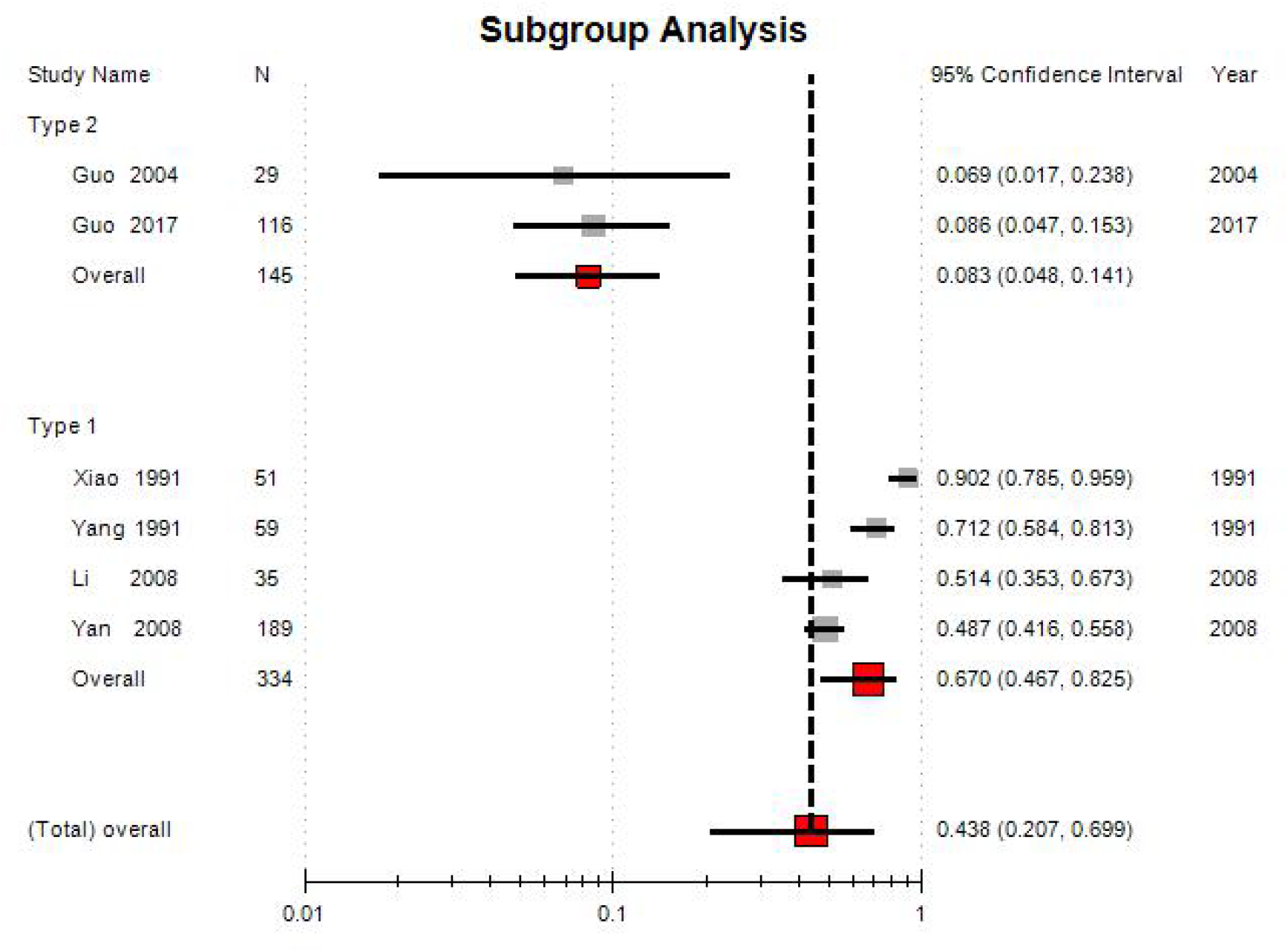
A Forest plot for proportion of the patients with phlegmonous appendicitis or more severe appendicitis between two subgroups

**Fig 2 B.**
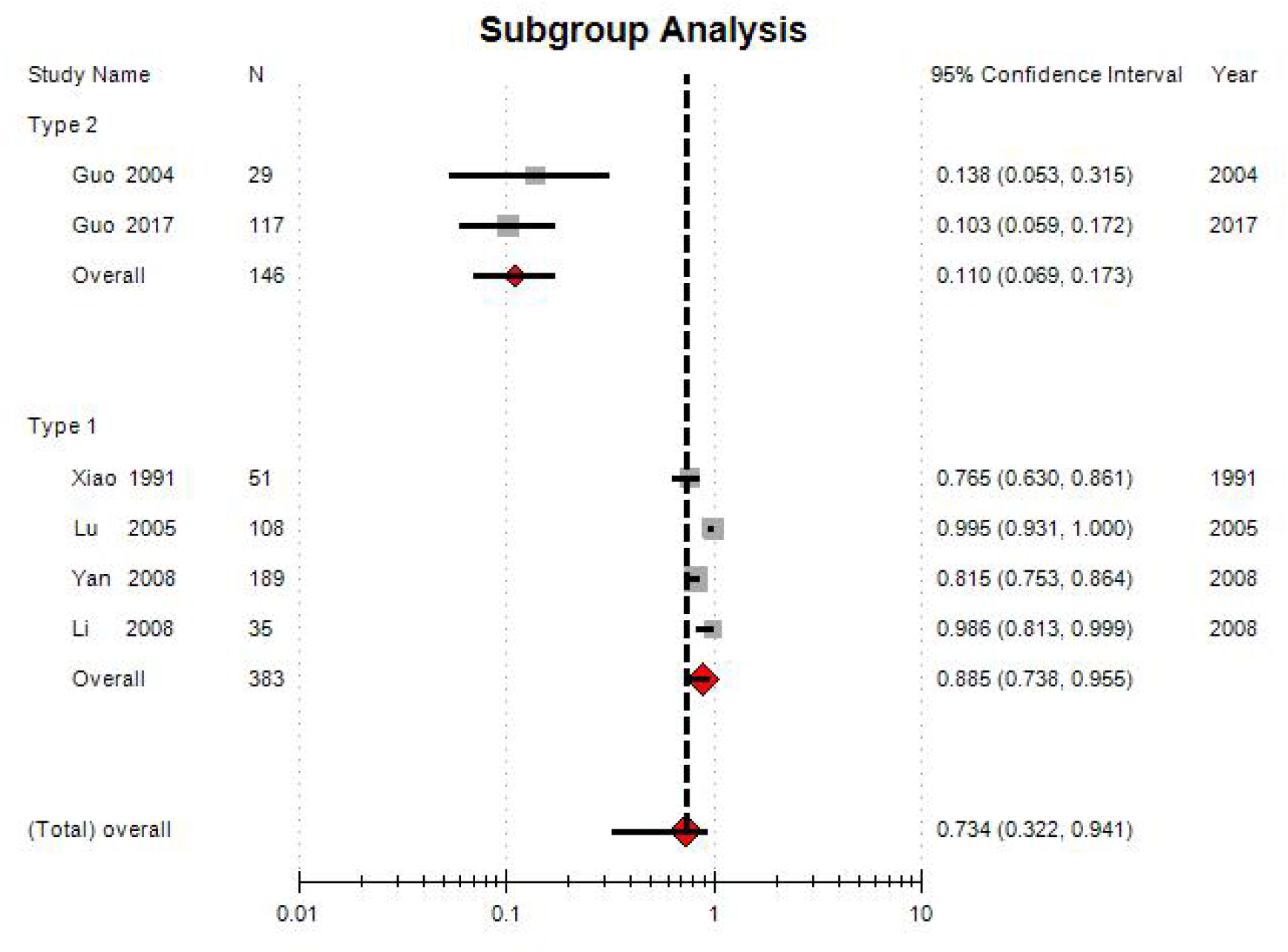
Forest plot for proportion of patients with elevated body temperature between Two subgroups

**Fig 2 C.**
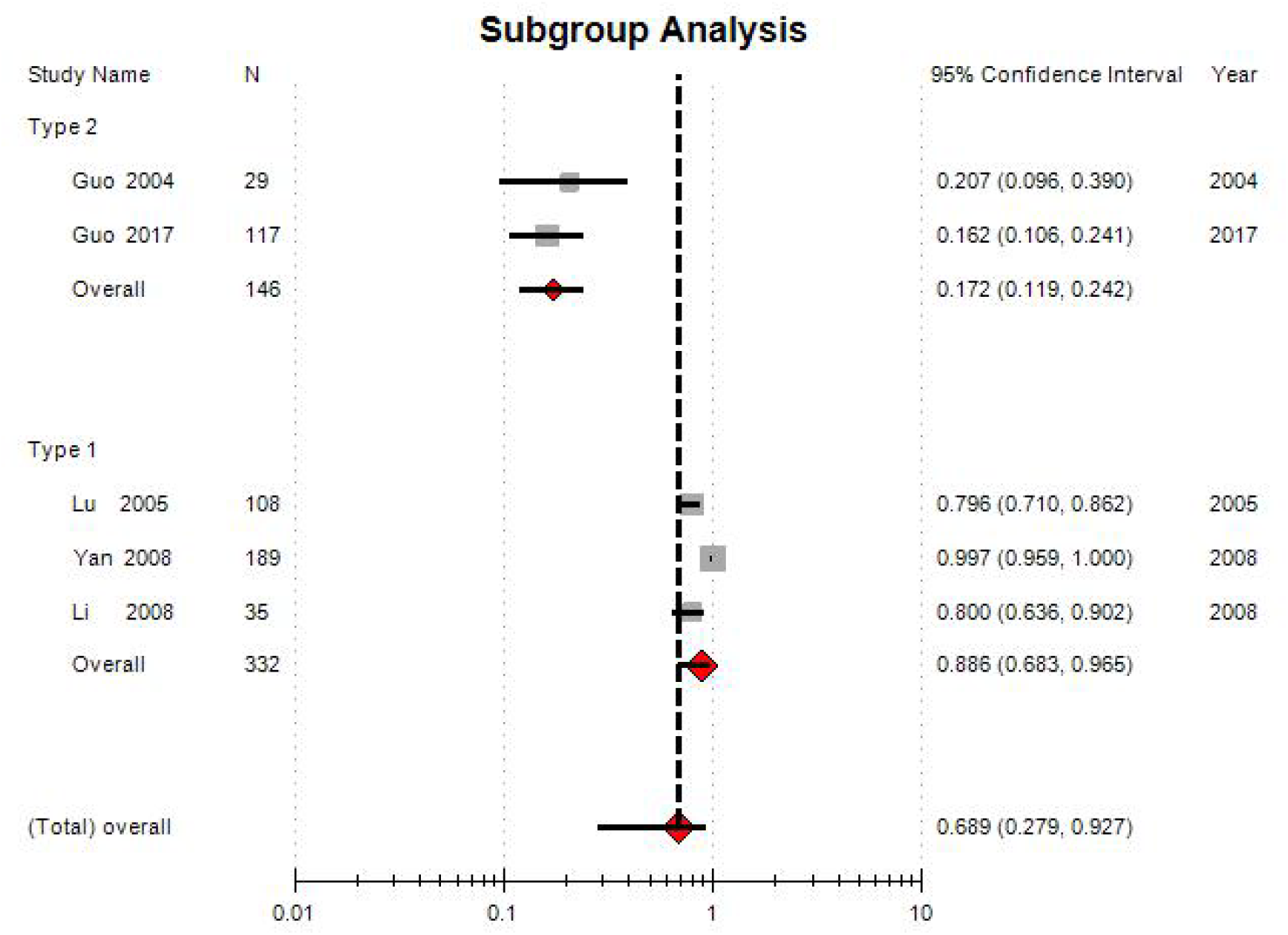
Forest plot for proportion of the patients with elevated white blood cells between two subgroups

**Fig 2 D.**
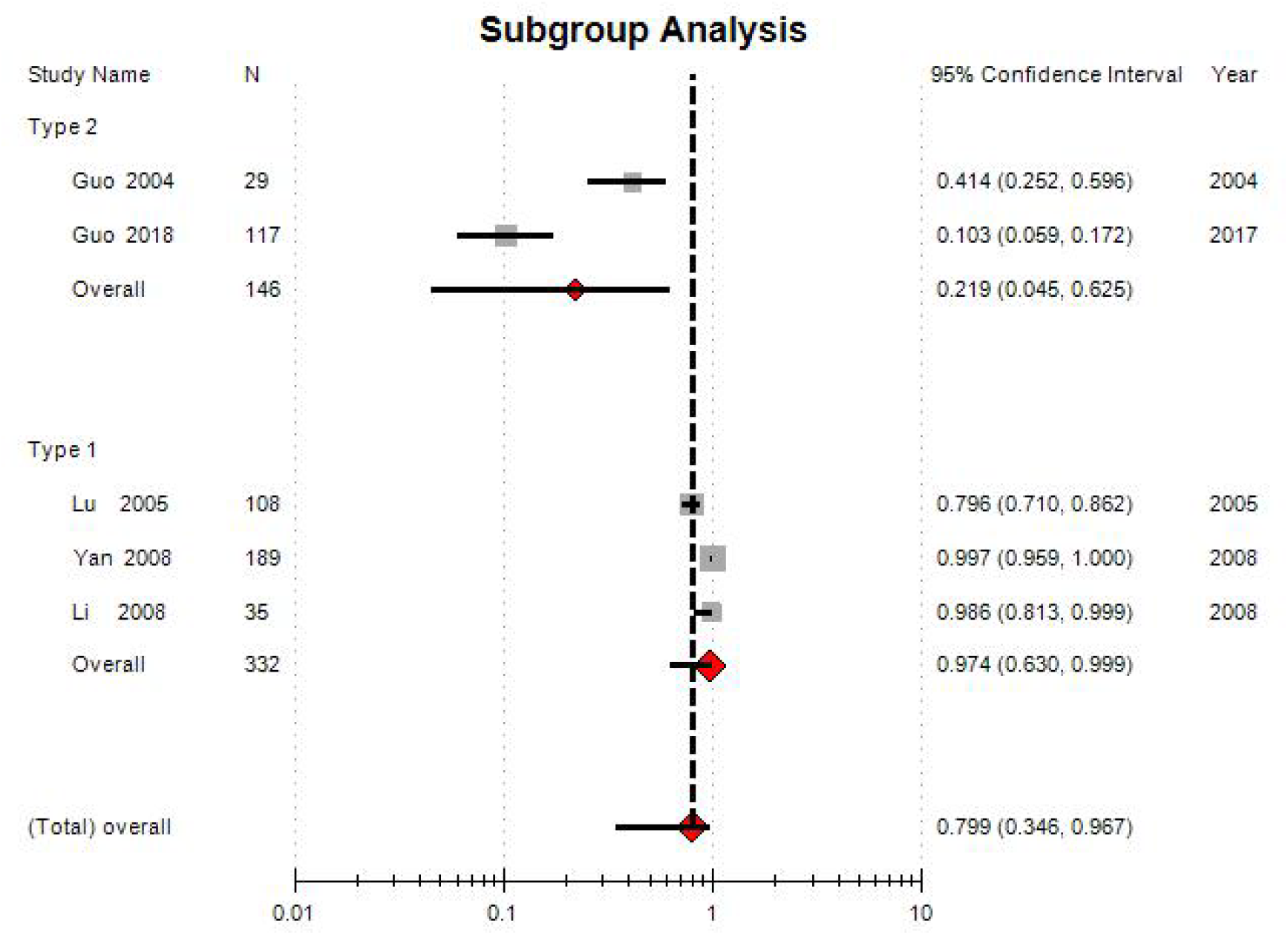
Forest plot for proportion of patients with elevated neutrophil percentage between two subgroups

Although we were not able to classify outbreak 7 into either of type 1 appendicitis and type 2 appendicitis because of no detailed clinical data available, however it showed that soldiers from minority were more susceptible.

According to imputation of average patients’ time of type 1 appendicitis, 27 hours were for non-perforated appendicitis and 41 hours for perforated appendicitis. However, for average patients’ time of type 2 appendicitis, outbreak 5 and outbreak 10 were about 50 hours and 112.8 hours respectively.

Epidemiological features, see table 1. All these clusters/outbreaks from reports occurred in group living units except outbreak 1 occurring in community, USA where most patients were students. Among them, 6 clusters/outbreaks occurred at schools and college, and the other 3 at camps in 7 provinces and autonomous regions. The incidence in female students were higher than that in male students (outbreak 5, outbreak 9 and outbreak 10), because female students contacted each other frequently and had similar living habit (outbreak 5 and outbreak 10); New students and new soldiers, especially from remote area and minority (outbreak 4, outbreak 5, outbreak 7, outbreak 9 and outbreak 10) are more susceptible; The new endemic focus of appendicitis can form and even persisted for several years or decades (outbreak 4, outbreak 8, outbreak 9 and outbreak 10).The potential transmission routes may included food-borne transmission (outbreak 1-3) and fomite transmission (outbreak 5 and outbreak 10).

Quality assessment. According to study-level assessment, one report met the basic requirement for outbreak introduced by Reingold and found infectious agent^37^. Another met the basic requirement of outbreaks introduced by Reingold, but did not detect infectious agent because of the limited conditions.^47^ Still another met the requirement partially, but the authors did not think of infectious etiology and so they did not investigate transmission route^42^. The rest of reports was case series of appendicitis from cluster/outbreak.^39–45^ Among them, only one report presented transmission route.^39^ However, according to outcome-level assessment, the outcome measure of these patients started as high-quality evidence. The reasons were as follows: In the GRADE approach, randomized trials start as high-quality evidence and observational studies as low-quality evidence. Because the outcome measures of these patients in cluster/outbreak did not need control group, there were not risk of bias in randomized trials and observational study. According to GRADE approach, case series can also provide high-quality evidence^63^. Considering that these patients were admitted into tertiary hospitals that is the first class hospital in China and the outcome measures of appendicitis were reliable. So we specified quality of evidence for the outcome measures as high, see table 2 and supplement 6.

**Table 2.**
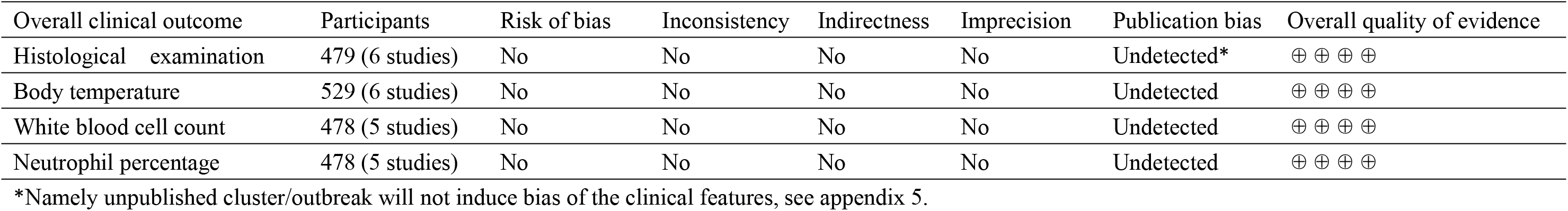
GRADE analysis: Overall clinical outcome of patients with appendicitis in cluster/outbreak―quality assessment

| Overall clinical outcome | Participants | Risk of bias | Inconsistency | Indirectness | Imprecision | Publication bias | Overall quality of evidence |
| --- | --- | --- | --- | --- | --- | --- | --- |
| Histological examination | 479 (6 studies) | No | No | No | No | Undetected* | ⊕ ⊕ ⊕ ⊕ |
| Body temperature | 529 (6 studies) | No | No | No | No | Undetected | ⊕ ⊕ ⊕ ⊕ |
| White blood cell count | 478 (5 studies) | No | No | No | No | Undetected | ⊕ ⊕ ⊕ ⊕ |
| Neutrophil percentage | 478 (5 studies) | No | No | No | No | Undetected | ⊕ ⊕ ⊕ ⊕ |

\*Namely unpublished cluster/outbreak will not induce bias of the clinical features, see appendix 5.

Sensitivity analysis showed that differences of outcome measures were not changed substantially between two types of appendicitis (Data not shown).

## Discussion

According to the definition of cluster/outbreak by CDC, the most reports included belonged to outbreaks. Our study demonstrated that cluster/outbreak of appendicitis occurred more often than expected. We have presented 10 outbreaks of appendicitis occurring in 7 provinces and autonomous regions in China, and one occurring in USA. As far as we know, this is the most detailed summarization of clusters/outbreaks of appendicitis. All clusters/outbreaks of appendicitis occurred in group living units except one occurred in community. The features of distribution will provide methods to find new cluster/outbreak. Because appendicitis is not endemic disease, our finding suggest that cluster/outbreak of appendicitis should also occur widely worldwide and can be found using same methods as we did in China. In fact, cluster/outbreak of appendicitis occurred more frequently than we realized in our systematic review. We can not report other schools where cluster/outbreak of appendicitis occurred, because these schools were not willing to collaborate with us.^47^

According to outcome-level assessment, our outcome measures are of high quality for data of clinical features and distribution features of patients in cluster/outbreak. Sensitivity analysis showed that differences of outcome measures were stable.

Our study may also provide more reliable method to differentiate different type of appendicitis before operation than analysis of secular trend and clinical data from sporadic patients.^4–7^ Through comparison of natural histories and clinical features, we demonstrated at least two types of independent entities, namely type 1 appendicitis and type 2 appendicitis. In future, more entities of appendicitis may be demonstrated through study of cluster/outbreak.

7 of 10 clusters/outbreaks were type 1 appendicitis. Their natural history showed continuum of acute simple appendicitis, acute phlegmonous appendicitis, gangrenous appendicitis, perforated appendicitis and so forth. The different forms of inflammatiom in appendices were histological features of different stage of same entity, not different entity. The clinical features showed elevated body temperature, WBC and NP in majority patients, but the patients’ time was shorter than type 2 appendicitis. Because majority of clusters/outbreaks belonged to type 1 appendicitis and its features similar to sporadic appendicitis, type 1 appendicitis likely represent most sporadic appendicitis and demonstrated classic description of natural history of appendicitis from non-perforated appendicitis to perforated appendicitis, which do not support modern classification that perforated appendicitis and non-perforated appendicitis are different entities.

2 of 10 clusters/outbreaks were type 2 appendicitis. Their natural history showed that most patients had acute simple appendicitis and only a few patients had phlegmonous appendicitis, and gangrenous appendicitis, therefore type 2 appendicitis belongs to non-perforated appendicitis. Most patients had normal body temperature, WBC and NP, and different histological features with hemorrhage and infiltration of eosinophils. This differs from sporadic non-perforated appendicitis indicating that type 2 appendicitis was a new entity of non-perforated appendicitis. The patients’ time was much longer than that of type 1 appendicitis. It means that is not reliable to early diagnose different type of appendicitis just through patients’ time.

Our study showed that current classification may misdiagnose different stage of same entity of appendicitis as two independent entities, namely perforated appendicitis and non-perforated appendicitis ^4–7^ or complex appendicitis and simple appendicitis.^8^ Therefore differences of clinical features between sporadic perforated appendicitis and sporadic non-perforated appendicitis are not due to different entities, but due to different stage of the same entity, namely differences of early stage of appendicitis and late stage. It can explain the reason why patients’ time of perforated appendicitis is longer than that of non-perforated appendicitis clinically. According to study of cluster/outbreak, we did not demonstrate that sporadic perforated appendicitis and sporadic non-perforated appendicitis are two independent entity as hypotheses described.^4–7^ Because if they are two independent entities, we should find such results, namely almost every patient in cluster/outbreak had either perforated appendicitis or classic non-perforated appendicitis.

Considering existence of type 2 appendicitis, we suggest to diagnose non-perforated appendicitis as type 2 appendicitis preliminarily if patient’s body temperature, WBC, and NP is normal, and have longer patient’s time than 50 hours. After appendectomy, diagnosis can be confirmed pathologically.

Our results provided more sufficient epidemiological evidence to support infectious etiology of appendicitis. For examples, the students and soldiers from remote areas, the minorities, and female students and the new comers are more susceptible than native students and soldiers.^37,40–47^ High attack rates in female students were associated with their living habit. New endemic location can form and persist for years and decades.^37,40,41, 44–47^ Transmission routes were associated with food borne transmission^35, 39^ and fomite transmission;^37,47^ Measures for control of infectious diseases seem to be effective to prevent appendicitis.^47^ Since 2009, new studies have provided compelling evidence of an association between appendicitis and the presence of Fusobacteria in the appendices,^64–69^ and Fusobacteria were also found in clustering patients we reported in 2012. ^38^

Study of cluster/outbreak will provide new methods to confirm causal association between microbiota and different entity of appendicitis. Jackson found that five taxa were increased in appendices in sporadic patients with perforated vs. non-perforated appendicitis: Bulleidia, Fusibacter, Prevotella, Porphyromonas, Dialister.^70^ As sporadic perforated appendicitis and non-perforated appendicitis may be different stage of the same entity, the increased five taxa may reflect the difference of microbiota between different stages. Because appendicitis is acute abdomen, the cluster/outbreak must occur if it is mainly communicable disease. Through studying patients in cluster/outbreak, the difference in microbiota between different entities of appendicitis may be confirmed. Further we may make etiological diagnosis for appendicitis and confirm proportion of communicable appendicitis and non-communicable appendicitis clinically, and in future, improve diagnosis and treatment of appendicitis.

## Limitation

Our reports were all case series except three of them which were studies.^37,46,47^ The case series did not described epidemiological features in detail and not conduce to confirming infectious etiology. Some reports did not provide data of elevated body temperature, WBC, NP and average patient’s time, and did not describe detailed histological features.^35,41,42^ Percentage of patients with elevated WBC in outbreak 9 and the average patients’ times for non-perforated appendicitis and perforated appendicitis of type 1 appendicitis were imputed based on references. We excluded some reports of clusters/outbreaks with no histological diagnoses, so there should be more clusters/outbreaks worldwide. As most cluster/outbreaks occurred in China, it mean that publication bias may exist, but main features of type 1 appendicitis and type 2 appendicitis will not be changed for it.

## Conclusion and suggestion for future work

We confirmed common settings of outbreak/cluster of appendicitis and provided new method to demonstrate different entities of appendicitis and causal association between between microbiota and different types of appendicitis. We did not demonstrate the current hypothesis that sporadic perforated appendicitis and sporadic non-perforated appendicitis may be different entities. Our epidemiological evidence supports infectious etiology of appendicitis. Future study should carry out surveillance or retrospective study for group living units to find new outbreak/cluster of appendicitis, further confirm different entities of appendicitis, causal association between infectious agents and appendicitis, and improve modern understanding from study of sporadic patients.

## Supporting information

manuscript

strobe checklist

## Author Contributors

Contributors: GY, TSY, GYT and LGZ designed the study. GY, GYT collected references. GY, GYT, TSY and LGZ analyzed and interpreted the data. GYT drafted the version of the manuscript.

All authors contributed and approved the manuscript. The corresponding authors attests that all listed authors meet authorship criteria and that no others meeting the criteria have been omitted.GY and TSY are the guarantors.

## Ethical approval

Not needed.

## Data sharing

No additional data available.

## Transparency

The lead author affirms that the manuscript is an honest, accurate, and transparent account of the study being reported; that no important aspects of the study have been omitted; and that any discrepancies from the study as planned (and, if relevant, registered) have been explained.

This is an Open Access article distributed in accordance with the Creative Commons Attribution Non Commercial (CC BY-NC 4.0) license, which permits others to distribute, remix, adapt, build upon this work non-commercially, and license their derivative works on different terms, provided the original work is properly cited and the use is noncommercial. See: http://creativecommons.org/licenses/by-nc/4.0/.

---

Fig 2 part A: 67% percent of the patients with type 1 appendicitis (I^2^=0.47) had phlegmonous or more severe appendicitis, which was 8.1 times as much as that (8.3%) of the patients with type 2 appendicitis (I^2^=000) (overall I^2^=0.49).

Fig 2 part B: 88.5% percent of the patients with type 1 appendicitis (I^2^=0.43) had elevated body temperature, which was 8.1 times as much as that (11%) of the patients with type 2 appendicitis (I^2^=0. 00) (overall I^2^=0.49).

Fig 2 part C: 88.6% percent of the patients with type 1 appendicitis (I^2^=0.45) had elevated WBC, which was 5.2 times as much as that (17.2%) of the patients with type 2 appendicitis (I^2^=0.00) (overall I^2^=0.46).

Fig 2 part D: 97.4% percent of the patients with type 1 appendicitis (I^2^=0.00) had elevated NP, which was 4.5 times as much as that (21.9%) of the patients with type 2 appendicitis (I^2^=0.48) (overall I^2^=0.49).

