## Supplementary material for "Demonstration of different entity of appendicitis and related causes of disease through study of cluster/outbreak: Systematic Review and Meta Analysis": manuscript

**Supplementary 1-6**

Supplementary 1. Search strategy: PUBMED

1. Acute appendicits. ti, ab
2. Cluster. ti, ab
3. Outbreak. ti, ab
4. 2 or 3
5. 1 and 4

25 papers of cluster/outbreak of acute appendicitis were retrieved.

Supplementary 2: Cluster: “an aggregation of cases of a disease, injury, or other health condition (particularly cancer and birth defects) in a circumscribed area during a particular period without regard to whether the number of cases is more than expected”. Outbreak: **“**the occurrence of more cases of disease, injury, or other health condition than expected in a given area or among a specific group of persons during a specific period. Usually, the cases are presumed to have a common cause or to be related to one another in some way” or “epidemic limited to localized increase in the incidence of disease”

Supplementary 3: According to PRIAMA suggestion^53^, missing data may be imputed from other information. For the percentage of patients with elevated WBC and NP, outbreak 8 did not present both and outbreak 9 did not present WBC. To avoid enlargement of difference between type 1 appendicitis and type 2 appendicitis, we imputed 80% as the the percentage based on the references^14,53^ , which is less than percentage presented in the references. For average patients’ time, except outbreak 5 and outbreak 10, the other reports present only shortest patients’ time and longest patients’ time with no average patients’ time. We selected three high quality and large sample papers as references.^9, 15, 17^ Among the three papers, the longest average patients’ time for nonperforated appendicitis was about 27 hours, for perforated appendicitis about 41 hours. Based on the results, we select 27 hours as average patients’ time for nonperforated appendicitis of the other reports and 41 hours as that for perforated appendicitis to avoid enlargement of difference of patients’ time between type 1 appendicitis and type 2 appendicitis.

Supplementary 4: High: We are very confident that the true effect lies close to that of the estimate of the effect; Moderate: We are moderately confident in the effect estimate: The true effect is likely to be close to the estimate of the effect, but there is a possibility that it is substantially different; Low: Our confidence in the effect estimate is limited: The true effect may be substantially different from the estimate of the effect; Very low: We have very little confidence in the effect estimate: The true effect is likely to be substantially different from the estimate of effect.

Supplementary 5: Fig 2 part A: 67% percent of the patients with type 1 appendicitis (I^2^=0.47) had phlegmonous or more severe appendicitis, which was 8.1 times as much as that (8.3%) of the patients with type 2 appendicitis (I^2^=000) (overall I^2^=0.49).

Fig 2 part B: 88.5% percent of the patients with type 1 appendicitis (I^2^=0.43) had elevated body temperature, which was 8.1 times as much as that (11%) of the patients with type 2 appendicitis (I^2^=0. 00) (overall I^2^=0.49)

Fig 2 part C: 88.6% percent of the patients with type 1 appendicitis (I^2^=0.45) had elevated WBC, which was 5.2 times as much as that (17.2%) of the patients with type 2 appendicitis (I^2^=0.00) (overall I^2^=0.46)

Fig 2 part D: 97.4% percent of the patients with type 1 appendicitis (I^2^=0.00) had elevated NP, which was 4.5 times as much as that (21.9%) of the patients with type 2 appendicitis (I^2^=0.48) (overall I^2^=0.49).

Supplementary 6: As described above, the outcome measure did not need control group, so there was no bias of risk of GRADE. About inconsistency, point estimate of outcome measures varied very little and confidence intervals (CIs) did not show overlap in type1 appendicitis and type 2 appendicitis respectively, and I^2^ for each outcome measure of type 1 appendicitis and type 2 appendicitis was less than 50% , so inconsistency was not substantial. About imprecision, sample size of patients in cluster/outbreak met optimal information size. There was no indirectness. For publication bias, according to our experience, cluster/outbreak of appendicitis was more often than expected. As it was not paid attention to, more clusters/outbreaks of appendicitis was not published. Further research may find new entities, but clinical features of type 1 and type 2 appendicitis were not influenced substantially, namely unpublished cluster/outbreak will not result in bias of the clinical features. Accordingly, We are very confident in the outcome measures.
