## Supplementary material for "Demonstration of different entity of appendicitis and related causes of disease through study of cluster/outbreak: Systematic Review and Meta Analysis": strobe checklist

Moose checklist for cluster/outbreak

Checklist item	Reported on page

Reporting of background should include

Problem definition	4

Hypothesis statement	5

Description of study outcome(s)	5

Type of exposure or intervention used	No exposure.

Type of study design	5, cluster/outbreak

Study population	5, patients in cluster

Reporting of search strategy should include

Qualifications of searchers (eg, librarians and investigators)	7

Search strategy, including time period included in the synthesis and keywords	5-6

Effort to include all available studies, including contact with authors	5,7

Databases and registries searched	5

Search software used, name and version, including special features used (eg, explosion)	No

Use of hand searching (eg, reference lists of obtained articles)	5

List of citations located and those excluded, including justification	6

Method of addressing articles published in languages other than English	5-6

Method of handling abstracts and unpublished studies	5

Description of any contact with authors	7

Reporting of methods should include

Description of relevance or appropriateness of studies assembled for assessing the hypothesis 8

to be tested

Rationale for the selection and coding of data (eg, sound clinical principles or convenience) 6, supplement 2.

Documentation of how data were classified and coded (eg, multiple raters, blinding, and	7

interrater reliability)

Assessment of confounding (eg, comparability of cases and controls in studies where	n/a*

appropriate)

Assessment of study quality, including blinding of quality assessors; stratification or regression	8

on possible predictors of study results

Assessment of heterogeneity	9-10

Description of statistical methods (eg, complete description of fixed or random effects models, 10

justification of whether the chosen models account for predictors of study results,

dose-response models, or cumulative meta-analysis) in sufficient detail to be replicated

Provision of appropriate tables and graphics	7

Reporting of results should include

Graphic summarizing individual study estimates and overall estimate	12, figure 2

Table giving descriptive information for each study included	10

Results of sensitivity testing (eg, subgroup analysis)	11, 13-14

Indication of statistical uncertainty of findings	12 ,imputation

Reporting of discussion should include

Quantitative assessment of bias (eg, publication bias)	17

Justification for exclusion (eg, exclusion of non–English-language citations)	18

Assessment of quality of included studies	14

Reporting of conclusions should include

Consideration of alternative explanations for observed results	n/a※

Generalization of the conclusions (ie, appropriate for the data presented and	14

within the domain of the literature review)

Guidelines for future research	18

Disclosure of funding source	19

* We did not compare case and control. ※ We did not find alternative explanations for our results.
